# Transcription factors use a unique combination of cofactors to potentiate different promoter-dependent steps in transcription

**DOI:** 10.1101/2022.10.25.513774

**Authors:** Charles C Bell, Laure Talarmain, Laura Scolamiero, Enid YN Lam, Ching-Seng Ang, Omer Gilan, Mark A Dawson

## Abstract

Transcription factors use DNA binding domains to recognise specific sequences and transactivation domains to recruit the cofactor proteins necessary for transcription. However, how specific cofactors contribute to transactivation at different genes remains unclear. Here, we couple Gal4-transactivation assays with comparative CRISPR-Cas9 screens to identify the cofactors required by nine different transcription factors and nine different core promoters in human cells. We classify cofactors as ubiquitous or specific, discover novel transcriptional co-dependencies and demonstrate that submodules within large co-activator complexes, such as the tail 2 and kinase modules of Mediator, facilitate transcriptional elongation. Rather than displaying discrete mechanisms of action, we discover that each TF requires a unique combination of cofactors, which influence its ability to potentiate distinct steps in the transcriptional process. Our findings help reconcile models of cofactor-promoter compatibility by demonstrating that transcription at different classes of promoters is constrained by either initiation or pause release. These differences dictate cofactor compatibility and the dynamic range of gene expression. Overall, our screens provide insight into TF-cofactor relationships and their ability to potentiate different steps in transcription at different classes of promoters.

## Introduction

Eukaryotic transcription factors (TFs) regulate specific gene expression programs by using their DNA binding domains to recognise particular DNA sequences, and their specialised transactivation domains (TADs) to recruit the proteins required for gene activation (*1, 2*). This transactivation process requires transcriptional cofactors (cofactors); a broad class of proteins that includes chromatin remodellers, histone modifying enzymes and other multi-subunit protein complexes that link TFs with the transcriptional machinery (*3*). Cofactors, generally do not demonstrate DNA sequence specificity and are instead recruited to specific loci by TFs. However, a systematic understanding of specific cofactor recruitment is lacking (*1, 4–6*). Similarly, how specific cofactors activate different types of promoters remains poorly understood. Whilst recent studies have reported a degree of cofactor compatibility between particular cofactors and core promoters, the leading model of ‘biochemical compatibility’ remains contested (*7–10*). Therefore, a mechanistic explanation for why certain cofactors demonstrate promoter preference is yet to be established.

Addressing these questions requires new strategies to link TFs and promoters with the cofactors they require for transcription. Structural approaches, which have been critical in characterising the DNA binding domains of TFs, are unable to provide insights into the often unstructured TADs (*5, 11*). Moreover, as transcription is such a complex process, traditional functional approaches are generally unable to deconvolute the influence of the various regulatory inputs, such as the activating TF, collaborating TFs, the core promoter or cellular context. Therefore, a more reductionistic approach is required, one in which the factors that contribute to transactivation are studied independently of other variables.

## Results

### Establishing a transcription factor-based screening system

To address this challenge, we developed a screening system consisting of two major components: (i) a DNA binding domain of Gal4 fused to a transactivation domain of interest and (ii) a reporter containing Gal4 binding motifs upstream of a minimal CMV promoter that drives expression of a fluorescent reporter (Figure S1A-B). This reductionistic design enabled us to specifically isolate how the transactivation domain alone alters the cofactors required for transcription. As we wanted to assess the contribution of all transcriptional regulators, including those required for cell survival, we used an unstable variant of GFP (*12*). With this reporter, we could detect complete loss of transcription upon disruption of the catalytic subunit of RNA polymerase 2 (POLR2A) (Figure S1C), demonstrating the ability to identify common-essential proteins necessary for transcription. To minimise the influence of additional regulatory inputs, we developed an isogenic, constant reporter cell line, by integrating the reporter into a Cas9 expressing K562 clone. Into this constant reporter cell line, we introduced Gal4 fused to the TADs of interest and confirmed a complete loss of fluorescent signal upon Gal4 KO, validating that we are measuring transactivator-dependent effects on gene expression (Figure S1D). To provide orthogonal validation, we also introduced our reporter construct into the endogenous AAVS safe harbour locus and validated the specific requirement of proteins identified in our subsequent genetic screens (Figure S1E).

We first decided to evaluate the cofactors required by nine different TFs with diverse and important functions in synthetic biology, development and disease. These include VP64, c-MYB (MYB), EWSR1 (EWS), p65 (NF-kB), P53, IRF1, PU.1, NOTCH and the glucocorticoid receptor (GR). The specific TAD region used was based on the largely unstructured regions of the TFs, enriched for acidic, proline and glutamine residues and previously shown to have activation potential (Figure S1B and Data S1). Some of the TFs used, such as VP64 and MYB, are known to be particularly dependent on specific cofactors. To validate that we can capture this specificity, we used the Gal4-VP64 and Gal4-MYB cell lines and confirmed that they were preferentially affected by loss of these selective cofactors (*13, 14*) (Figure S1C).

Using these nine reporter cell lines, we performed CRISPR-Cas9 screens with a bespoke guide library targeting 1137 transcriptional regulators and chromatin associated proteins (Data S2). All of the screens were performed in parallel and samples were harvested at three timepoints (D5, D6, D7) to minimise technical variability and permit quantitative comparisons of the effects of different cofactors on TAD activity (Figure S2A-B, Methods). The screens identified a total of 239 out of 1137 genes in the library as required for at least one of the nine TFs (Data S3). As expected, the hits cluster closely together by STRING analysis, with clear enrichment of clusters involved in RNA polymerase initiation and elongation, the Mediator complex, SWI/SNF components and SET/COMPASS family members (Figure S2C). To ensure that we could separate transcriptional effects from effects on cell viability, we intersected the screen hits with dropout data from matched samples. Notably, approximately 30% of the genes required for cell growth were not required for transcription in any of the screens (Figure S2D). Similarly, integration with the cancer dependency map (DEPMAP) database, revealed that a large number of common-essential genes involved in chromatin regulation were not required for transactivation in any of our screens (Figure S2E). Together this demonstrates that our screens can identify essential transcriptional regulators and divorce their specific contribution to transcriptional activation from their requirement for cell viability.

### An overview of the specificity and heterogeneity of transcription factor-cofactor interactions

To provide a global overview of the screen results, we created a visualisation chart that shows the requirement for a range of key transcriptional regulators across the 9 TFs (Figure 1). Upon initial inspection, it is clear that our screens identified various proteins that are known to be directly involved in transcription. For example, the entire RNA polymerase 2 complex is required for all of the TFs (99/99 possible enrichments). Likewise, most components of the preinitiation complex, such as TFIIA, TFIIB, TFIIE, TFIIF, TFIIH, as well as the FACT complex and DSIF components appear to be ubiquitously required. Beyond these core transcriptional proteins, other coactivator complexes display more interesting patterns. For example, the Mediator complex, which bridges TFs with the core transcriptional machinery, has some subunits that are equally required for all TFs, while others display a large degree of heterogeneity. Some of these subunits are required together for the same specific TFs. For example, MED16, MED23, MED24 and MED25 are heterogeneously required, but have similar patterns of requirement, suggesting that these subunits are co-dependent and may function together to regulate transcription. Other more distal coactivators such as the SET/COMPASS complex, the Integrator complex, chromatin remodellers and transcriptional elongation components also appear to display a large degree of heterogeneity and interesting patterns of co-dependency.

**Figure 1).**
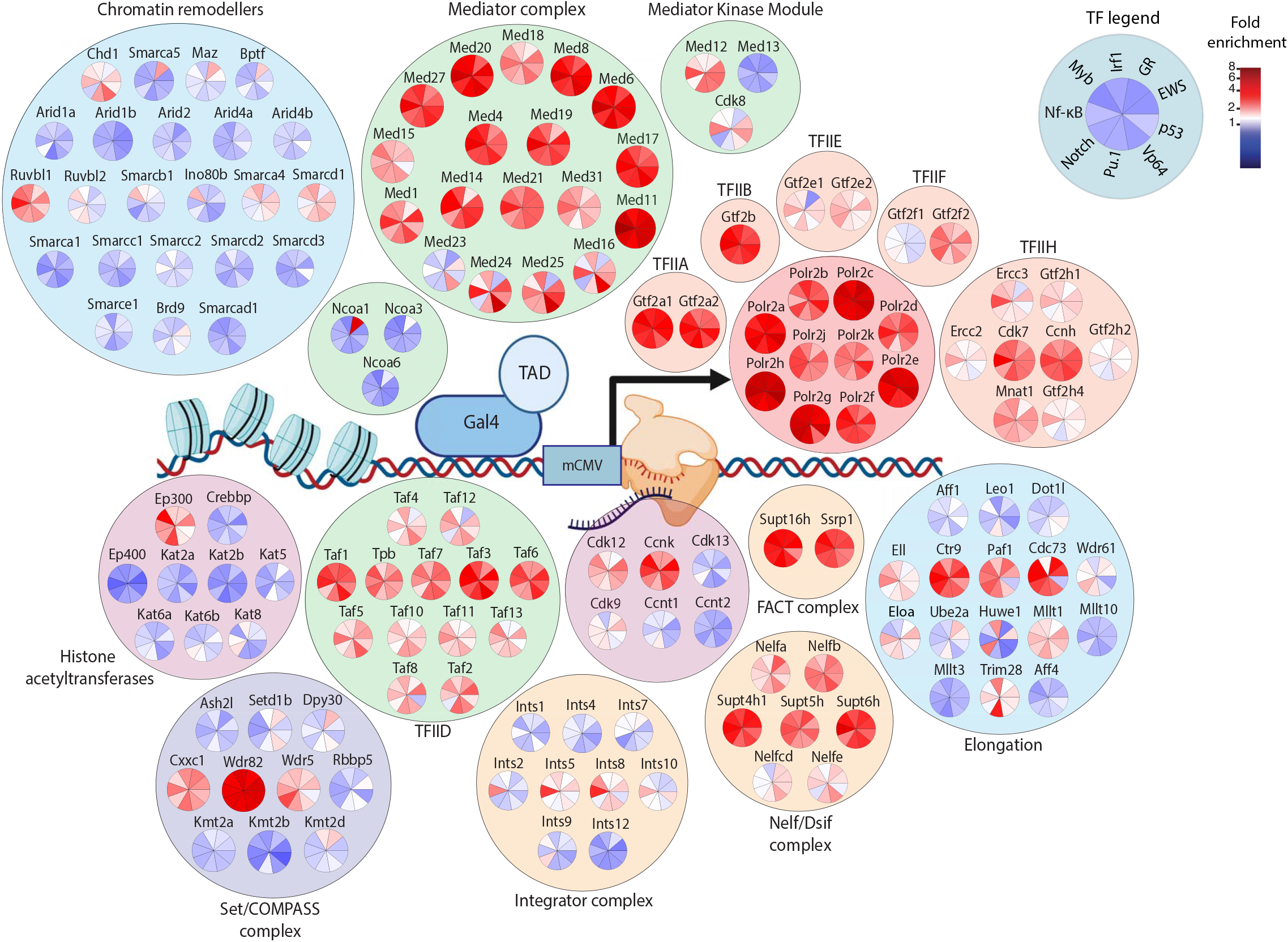
Comparative CRISPR screens identify the cofactors required for transactivation by nine different TFs. Visualisation chart demonstrating the requirement of key transcriptional cofactors for each of the nine transcription factors. The colour in each wedge reflects the fold enrichment score for each TF. Cofactors are organised into particular complexes based on their known complex associations or known molecular functions.

Reassuringly, our approach unbiasedly identified several previously reported interactions between TFs and cofactors, including MYB-EP300 (*13, 15*), MYB-TAF12 (*16*), VP64-MED25 (*14*), P53-CDK8 (*17*) and NOTCH-WDR5 (*18*) (Figure 1). In addition to these known interactions, several novel TF-cofactor associations were observed. One that was particularly striking was INTS5, which along with two other subunits of the Integrator complex, INTS2 and INTS8, was especially important for NF-kB transactivation (Figure 1, Figure S3A). To confirm that INTS5 is preferentially required for the endogenous activity of NF-κB, we treated K562 cells with TNF-α to stimulate NF-κB activity and assessed the effects in control and INTS5 KO cells. By both cell surface expression and global ChIP-seq analyses, we confirmed that INTS5 loss has a preferential effect on endogenous NF-kB activity (Figure S3). Therefore, our method was able to identify a submodule of the Integrator complex containing INTS5 (*19*) as critical for NF-kB activity, demonstrating the ability for our dataset to identify new TF-cofactor associations that reflect endogenous mechanisms of transactivation.

### Exploring the specificity and heterogeneity of cofactors involved in transactivation

Our comparative screens act not only as an important resource, but also provide the opportunity to obtain systematic insights into the transactivation process. To begin with, we explored the relationship between the specificity of a cofactor and its quantitative contribution to transcription. In general, we observed that cofactors that are broadly required by most TFs tend to have a larger effect on transcription (Figure 2A). Interestingly, we observed very few examples of cofactors with a potent and highly selective requirement, with the notable exception of NCOA1, which is a major dependency for GR-mediated transactivation. This suggests that, in general, TFs do not to have dedicated cofactors that contribute strongly to transcription.

**Figure 2).**
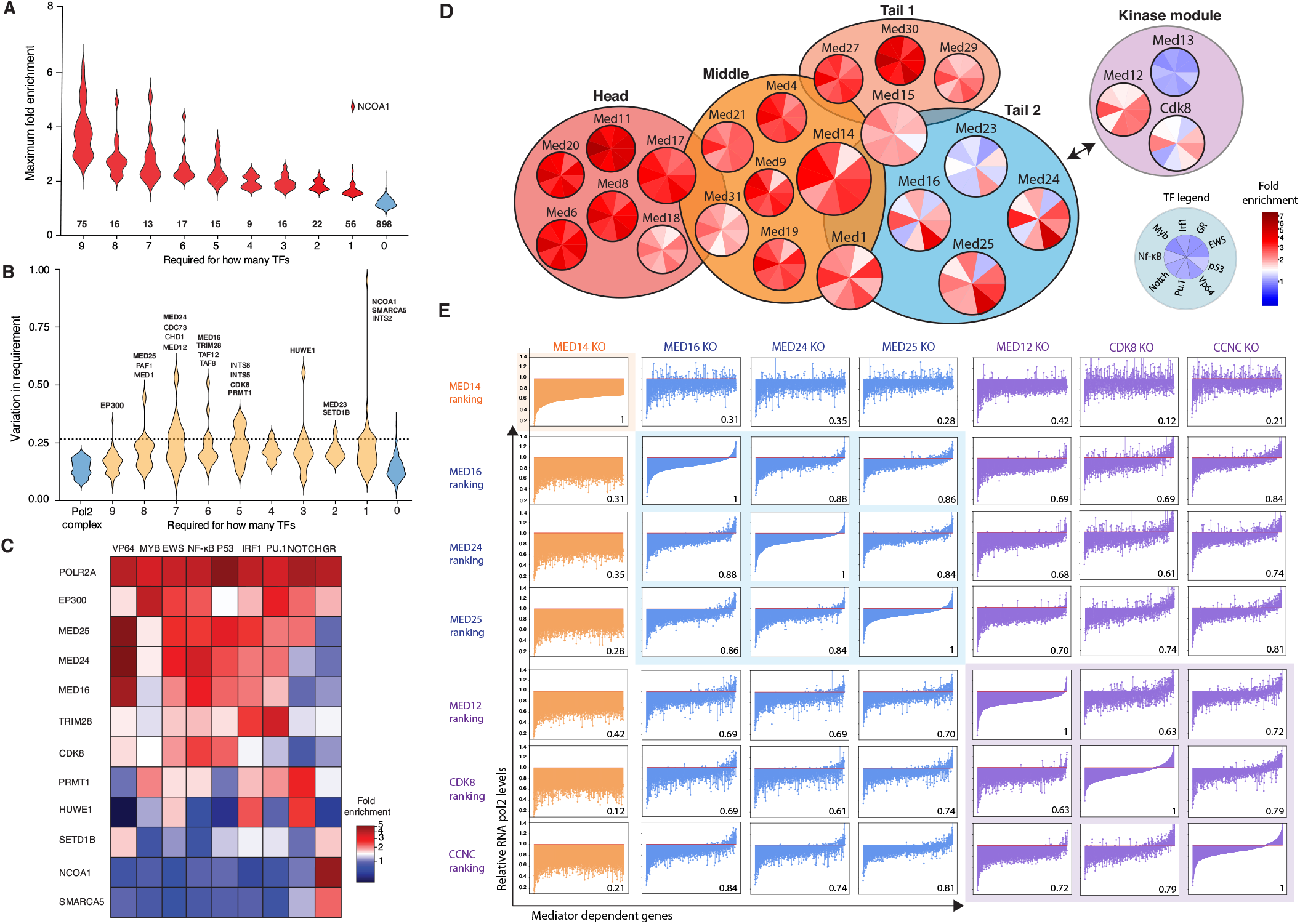
Transcription factors display a diverse range of activation mechanisms. **(A)** Violin plot of maximum fold enrichment score for each gene in the library across all 9 screens. The number of genes in each category is displayed. **(B)** Violin plot of the degree of variability for each gene across the 9 screens. Variability calculated by dividing the standard deviation for each hit across the nine TFs by the average fold enrichment across the nine TFs. Selected variable genes in each category are displayed. Genes that were independently validated are highlighted in bold. **(C)** Heatmap of fold enrichment score for selected heterogeneous hits that have been independently validated by individual KO. **(D)** Visualisation chart of the requirement of Mediator complex subunits across the nine TFs. **(E)** Correlation matrix of change in RNA polymerase 2 levels at Mediator dependent genes upon individual subunit KO. The genes are ordered based on the degree of reduction in each KO sample. Spearman correlation between the rank order is displayed.

A lack of potent, selective cofactors does not necessarily imply a generic mechanism for TF transactivation. Instead, our data suggests that rather than using dedicated cofactors, TFs may achieve specificity by using a unique combination of cofactors (Figure 1). To isolate the heterogeneously required cofactors, we calculated a variability metric for each gene across the 9 screens (detailed in Methods). We anchored this analysis by comparison with genes that are not expected to be variable, such as the RNA polymerase 2 complex. This identified ~100 cofactors that display heterogeneity in their requirement, some of which are required for a large number of TFs, such as EP300 or CHD1, and others that are required by few TFs, such as SETD1B or NCOA1 (Figure 2B). Interestingly, we observed that even broadly required cofactors, such as EP300, can display variable contributions to transactivation. To provide confidence in these results, we selected eleven of these heterogeneously required cofactors, and independently quantified their contribution to transactivation by each TAD (Figure S4A-B). Overall, we observed a high degree of concordance between the enrichment scores reported by the screen and the reduction in GFP signal in each respective TAD line, demonstrating that the heterogeneity is genuine and reinforcing the highly quantitative nature of the screens (Figure S4A-B).

Overall, our data suggests that there are no simple rules to explain when specific individual cofactors are required. This can be observed through the absence of clear patterns of requirement across the different TFs (Figure 1, Figure 2C). For instance, PU.1, is highly dependent on EP300, TRIM28 and PRMT1. Whilst NOTCH, which also requires EP300 and PRMT1, does not require TRIM28 (Figure 2C). Similarly, although EWS and VP64 both require CDK8 and TRIM28; EWS requires HUWE1 whereas VP64 does not (Figure 2C). These examples illustrate the more general observation (Figure 1) that that each TF uses a unique set of cofactors to activate transcription.

### Mediator tail 2 and the kinase module often function together to regulate transactivation

While most cofactors do not fall into discrete categories, some cofactors do display highly correlated patterns of requirement (Figure 1). By looking for hits with similar patterns of requirement across the 9 different TFs, we confirmed that in general there is a poor correlation between most of the heterogeneously required cofactors (Figure S4C). However, this codependency analysis did identify certain clusters of highly correlated, co-dependent cofactors, particularly within multi-subunit co-activating complexes. These include tail 2 of the mediator complex (MED16, MED23, MED24, MED25), CDK12 and CCNK, subunits of the Integrator complex (INTS2, INTS5, INTS8) and PAF1/WDR61 (Figure S4D).

Due to its role directly linking TFs with the RNA Pol II complex, we were particularly interested in the co-dependencies observed within the Mediator complex. The mammalian Mediator is organised into 5 major modules – the head, middle, tail 1, tail 2 and the sub-stoichiometric kinase module (*20*, *21*) (Figure 2D). Whilst the structural composition of Mediator has been well characterised, the functional relationships between subunits within the complex remain poorly understood. Our data suggested that the head, middle and tail 1 modules are ubiquitously required for transactivation (Figure 2D). Interestingly, most of the functional heterogeneity within Mediator exists within the tail 2 module, the kinase module and MED1, which is located structurally proximal to tail 2 (Figure 2D).

We noted that the requirement for tail 2 is also well correlated with requirement for the sub-stoichiometric Mediator kinase module (Figure 2D, Figure S4C). VP64, EWS, NF-kB and P53 are highly dependent on tail 2 and the kinase module, while MYB and GR are largely tail 2 independent and show little to no requirement for any of the kinase module subunits (Figure 2C, Figure S4E). To test whether this co-dependency extends beyond our screening system, we performed KO experiments for a number of the tail 2 subunits (MED16, MED24, MED25), the kinase module (CDK8, MED12, CCNC) and a core structural subunit (MED14) and assessed whether the effects were correlated at endogenous genes. As expected, loss of the core subunit (MED14) led to a marked and relatively uniform decrease in RNA Pol II levels (Figure 2E). In contrast, loss of tail 2 subunits (MED16, MED24, MED25) or the kinase module (CDK8, MED12, CCNC) resulted in disproportionate effects on particular subsets of genes (Figure 2E). Importantly, the effects of disrupting the individual tail 2 and kinase subunits were highly correlated, with a clear concordance between the rank order of genes effected by MED16, MED24, MED25, MED12, CDK8 and CCNC loss (Figure 2E). Consistent with our screens, the correlation is strongest within tail 2, with a minority of tail 2 dependent genes, not dependent on CDK8 and CCNC (Figure 2E). Importantly, the co-dependency is highly specific to tail 2 and the kinase module, as there was little correlation with disruption of the core mediator subunit (MED14 KO) (Figure 2E). Taken together, our comparative screens uncover a previously uncharacterised functional association within tail 2, and between tail 2 and the kinase module of the Mediator complex, which likely has important implications for how particular TFs activate their targets.

### TFs use different cofactors to potentiate different steps in the transcription process

As transcription is a multistep process that involves promoter opening, initiation, pausing, elongation and termination, we hypothesised that TFs may recruit a diverse array of cofactors in order to influence different stages of transcription. To explore this hypothesis, we used ChIP-nexus to precisely map the location of RNA Pol II on the reporter construct in each of the GAL4-TAD cell lines (*22*). Remarkably, when the reporter is activated by different TFs, we observed a striking difference in the amount of Pol II within the gene body, relative to the amount at the pause site (Figure 3A). Therefore, TFs have different capacities to facilitate Pol II initiation and elongation, which due to our reductionistic design, can be attributed principally to their TADs and the different cofactors they recruit.

**Figure 3).**
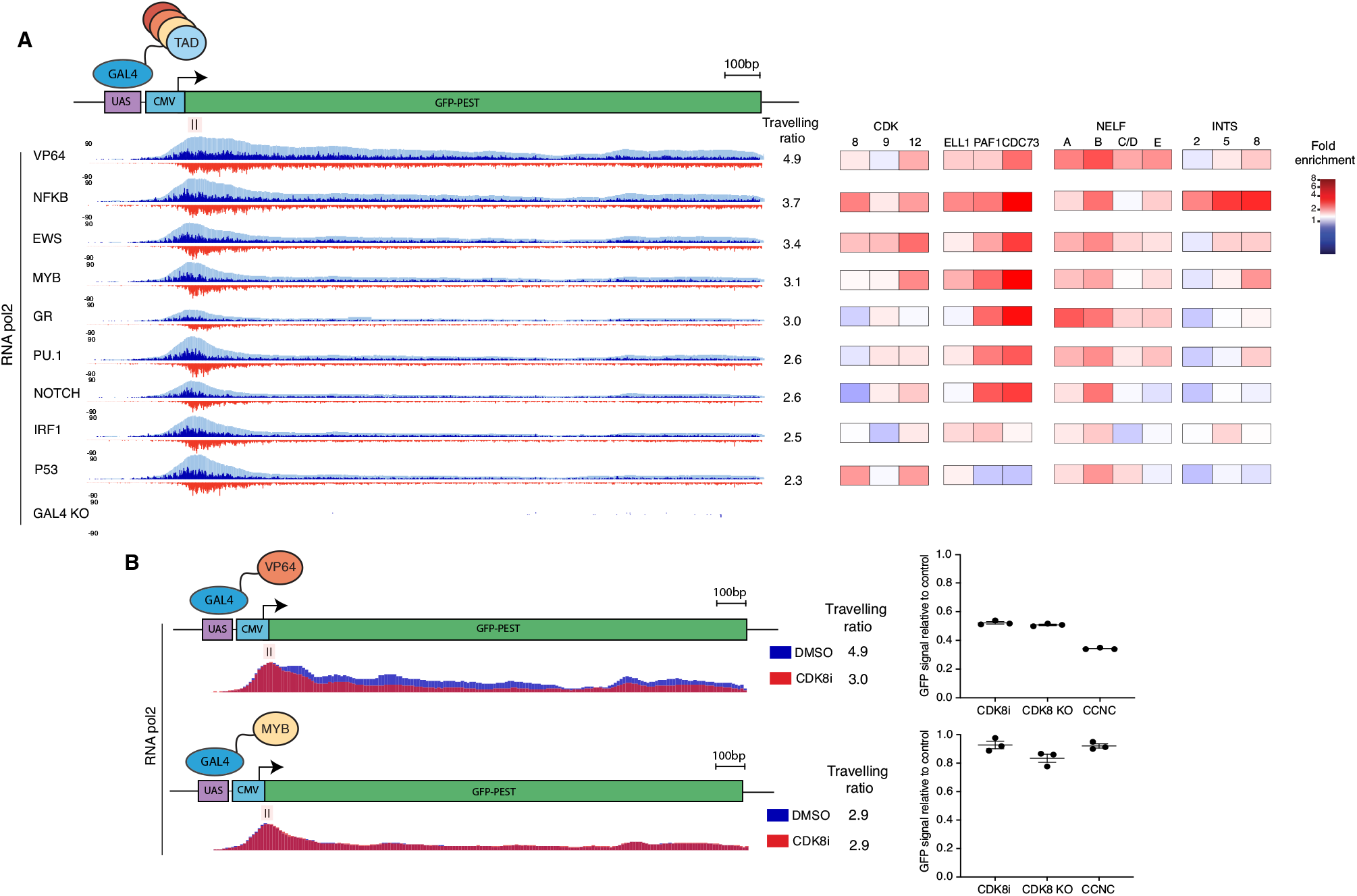
Transcription factors use different cofactors to trigger different steps in transcription. **(A)** RNA polymerase 2 ChIP-nexus coverage across the reporter construct in each of the GAL4-TAD cell lines. The travelling ratio and requirement for pause release cofactors by each TF is also displayed. Travelling ratio is calculated by dividing the total promoter signal by the total gene body signal. **(B)** RNA polymerase 2 ChIP-nexus coverage across the reporter construct in the Gal4-VP64 and GAL4-MYB cell lines treated for 1hr with DMSO or CDK8i. The transcriptional response to CDK8i, CDK8 KO and CCNC KO in these two cell lines is also shown.

Consistent with this idea, the ability to facilitate Pol II elongation correlated well with the requirement for cofactors associated with Pol II pause release (Figure 3A). Notably, the two TFs with the highest degree of pause release, VP64 and NF-kB, were the most dependent on the major pause release regulators including NELF and Integrator (Figure 3A). Interestingly, despite the fact that both of these TFs are strong activators of elongation, they are each most dependent on different regulators of pausing (Figure 3A), providing further support for the prospect of multiple, independent pause release checkpoints (*23*). We also observed that the requirement for positive regulators of transcriptional elongation, such as CDK8, CDK9, CDK12, ELL1, PAF1 and CDC73, corresponded with the travelling ratio for each transcription factor (Figure 3A).

While no cofactor alone is predictive of the ability to activate elongation, we were struck by the correspondence between the travelling ratio and the requirement for the Mediator kinase, CDK8. As we had previously identified the functional co-dependency between the CDK8 kinase module and tail 2 of the Mediator complex (Figure 2C-D), we considered the prospect that some TFs, such as VP64, interact with subunits in tail 2 to elicit the kinase module to potentiate elongation (*14*). To test this idea, we assessed the effects on Pol II in the VP64 cell line treated with a CDK8 inhibitor. These data showed that CDK8 inhibition substantially reduced the ability of the VP64-TAD to potentiate elongation (Figure 3B). In contrast, Pol II was unperturbed by CDK8 inhibition in cells expressing the MYB-TAD (Figure 3B), which is not reliant on CDK8 or tail 2 for transactivation (Figure 1). Together these findings suggest that these submodules of Mediator are specifically used by certain TFs to facilitate transcriptional elongation.

### Different submodules of TFIID are utilised at different core promoters

The transactivation process requires the cofactors recruited by TFs to converge onto a promoter where RNA pol II is loaded. Our screens identified that TFs recruit diverse sets of cofactors to potentiate different steps in transcription. However, it remains unclear how different promoters interact with these specific cofactors to influence the mechanisms of transactivation. To address this issue, we adapted our screening system to alter the core promoter of the reporter and maintain a constant transactivation domain. We chose to study several well-characterised core promoter elements: (i) the TATA box, (ii) a TATA-like element with reduced affinity for TBP (*24–26*), (iii) the Initiator sequence (Inr) and (iv) the polypyrimidine Initiator (TCT) sequence. Many of the genes that contain a TATA box have focussed promoters that are tissue specific, have a large dynamic range in gene expression and can be rapidly induced (*27, 28*). In contrast, the TCT element is present in housekeeping genes, in particular those that code for ribosomal proteins (*29*). These genes are often widely expressed across all tissues and have a narrow dynamic range in gene expression (*27*). Interestingly, previous reports have demonstrated that these different classes of promoter are differentially responsive to the NF-kB-TAD, suggesting that they have inherently different mechanisms of regulation (*30*). To further address this observation, we initially characterised the ability of NF-kB to activate transcription of these different classes of core promoters (Figure S5A). We chose to evaluate five core promoters containing a TATA box and Inr element, two core promoters containing a TATA-like sequence and Inr and two core promoters containing a TCT element (Figure S5A-B). We found that all the TATA and TATA-like core promoters containing an Inr element were compatible with transcription being strongly induced by the NF-kB TAD; in contrast the TCT promoters were activated to a much lower extent, indicating that they are incompatible (Figure S5C-D).

We next sought to identify the underlying mechanistic basis for the differential responsiveness/compatibility of these promoters to the NF-kB-TAD by creating nine independent reporter lines for each of these promoters and performing comparative cofactor screens using the same experimental design described earlier (Figure 1 and Methods). The results of these screens are summarised with another visualisation chart, which highlights the influence of the core promoter on the mechanisms of transcription (Figure 4A, Data S4). These data demonstrate a general requirement for the RNA Pol II complex and the components of the pre-initiation complex (TFIIA, TFIIB, TFIID, TFIIE, TFIIF and TFIIH). Interestingly however, we identified a differential requirement for the various submodules of TFIID with different core promoters. As expected, TBP is a major requirement for the activation of promoters with a TATA box (Figure 4A). Consistent with recent structural studies, in which TAF11 and TAF13 form a bridge linking TBP to TFIID (*31*), we identify that these two components of TFIID are required together with TBP. Notably, this submodule of TFIID is less required for promoters with a TATA-like element with lower affinity for TBP and is largely not required for the TCT-containing core promoters (Figure 4A). At TCT-containing promoters, we instead identify a major requirement for TAF1, TAF2, TAF7 and TAF8, which are structurally co-located and interact with promoter elements in a manner distinct from TBP (Figure 4A) (*31–33*). Together these data illustrate that TFIID-mediated assembly of the preinitiation complex is required across all of these core promoters, however different submodules of TFIID are required for the activation of core promoters that contain TATA and Inr elements, as compared to promoters with a TCT element (Figure 4A-B).

**Figure 4).**
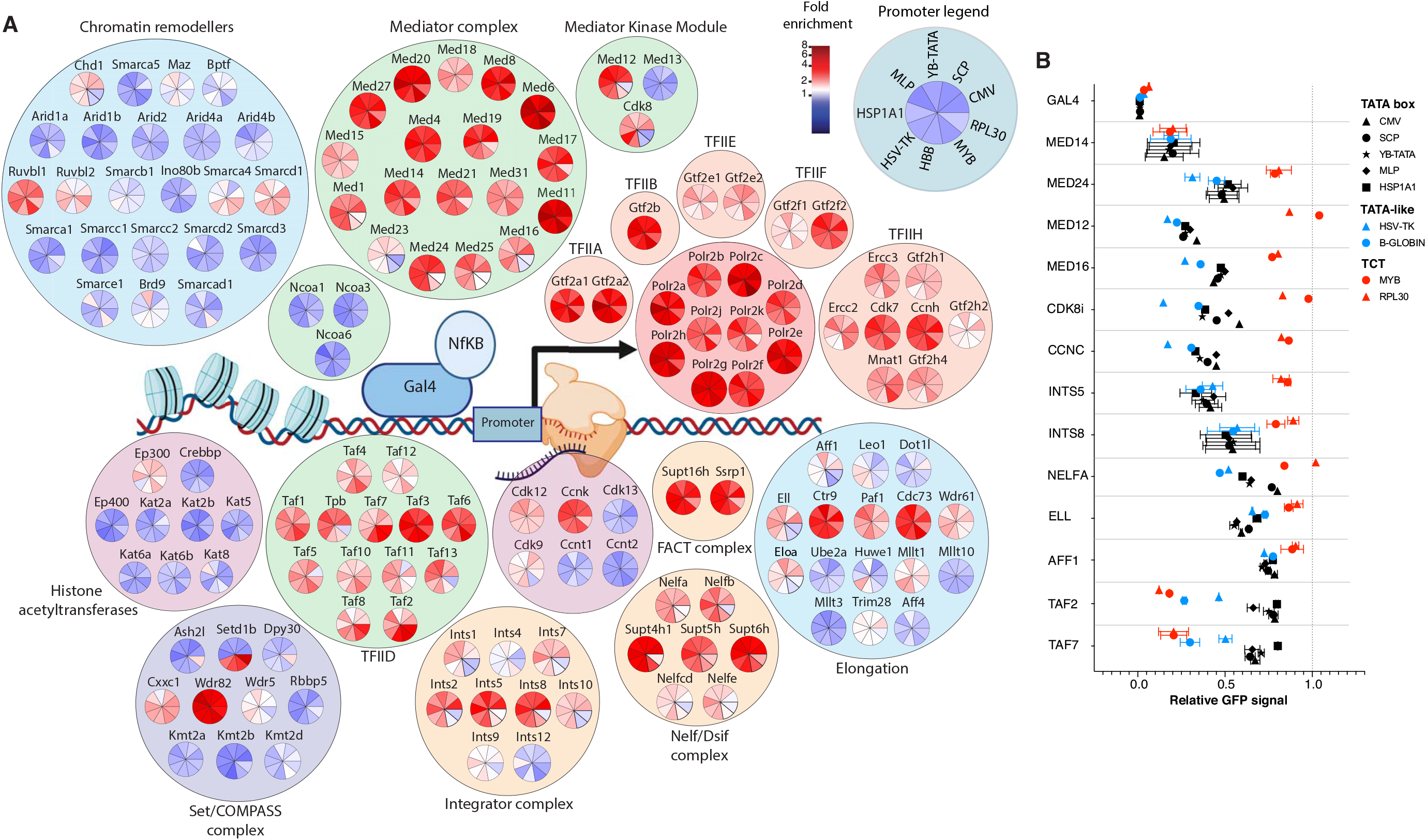
Comparative screens with 9 core promoters reveals a different mechanism of initiation and lack of requirement for pause release cofactors at incompatible promoters. **(A)** Visualisation chart demonstrating the requirement of key transcriptional cofactors for each of the nine core promotors activated by Gal4-NF-kB. The colour in each wedge reflects the fold enrichment score for each core promoter. Cofactors are organised into particular complexes based on their known complex associations or known molecular functions. For ease of comparison, the same cofactors are shown as Figure 1. **(B)** Validation of screen data through quantification of GFP signal. Quantification value is based on M.F.I and is compared to a SAFE guide control at D5 after infection with relevant sgRNAs. Most genes were validated with at least 2 sgRNAs, error bars reflect S.E.M between two sgRNAs.

### Identifying the mechanistic basis of cofactor-promoter incompatibility

Our previous results had identified that the TAD of NF-kB requires submodules of Mediator (Figure 2D-E), Integrator (Figure S3) and several other cofactors implicated in regulating the pause-release checkpoint (Figure 1 and Figure 3A). Interestingly, we observed that many of these cofactors, including the co-dependent tail 2 and kinase module of Mediator, the Integrator complex, NELF, DSIF, ELL, AFF1 and ELOA, are required at the compatible TATA and Inr promoters, but not required in the context of the incompatible TCT promoters (Figure 4A-B).

Based on these results, we hypothesized that cofactor-promoter incompatibility may arise because different promoter classes have distinct rate-limiting steps to transcription. To formally test this hypothesis, we performed RNA Pol II ChIP-nexus on our reporter constructs containing the compatible (TATA) and incompatible (TCT) promoters activated by NF-kB. The compatible TATA promoters displayed clear evidence of Pol II accumulation at the pause site, suggesting that pausing is the rate-limiting step (Figure 5A). In contrast, neither of the incompatible TCT promoters display detectable accumulation of Pol II around the TSS (Figure 5A). This suggests that cofactor/promoter incompatibility occurs when TFs recruit pause release cofactors to promoters at which pause-release is not the rate-limiting step. In line with these findings TATA, TATA-like and Inr elements are frequently found in genes where pause release plays a major role transcriptional regulation (*34–36*).

**Figure 5).**
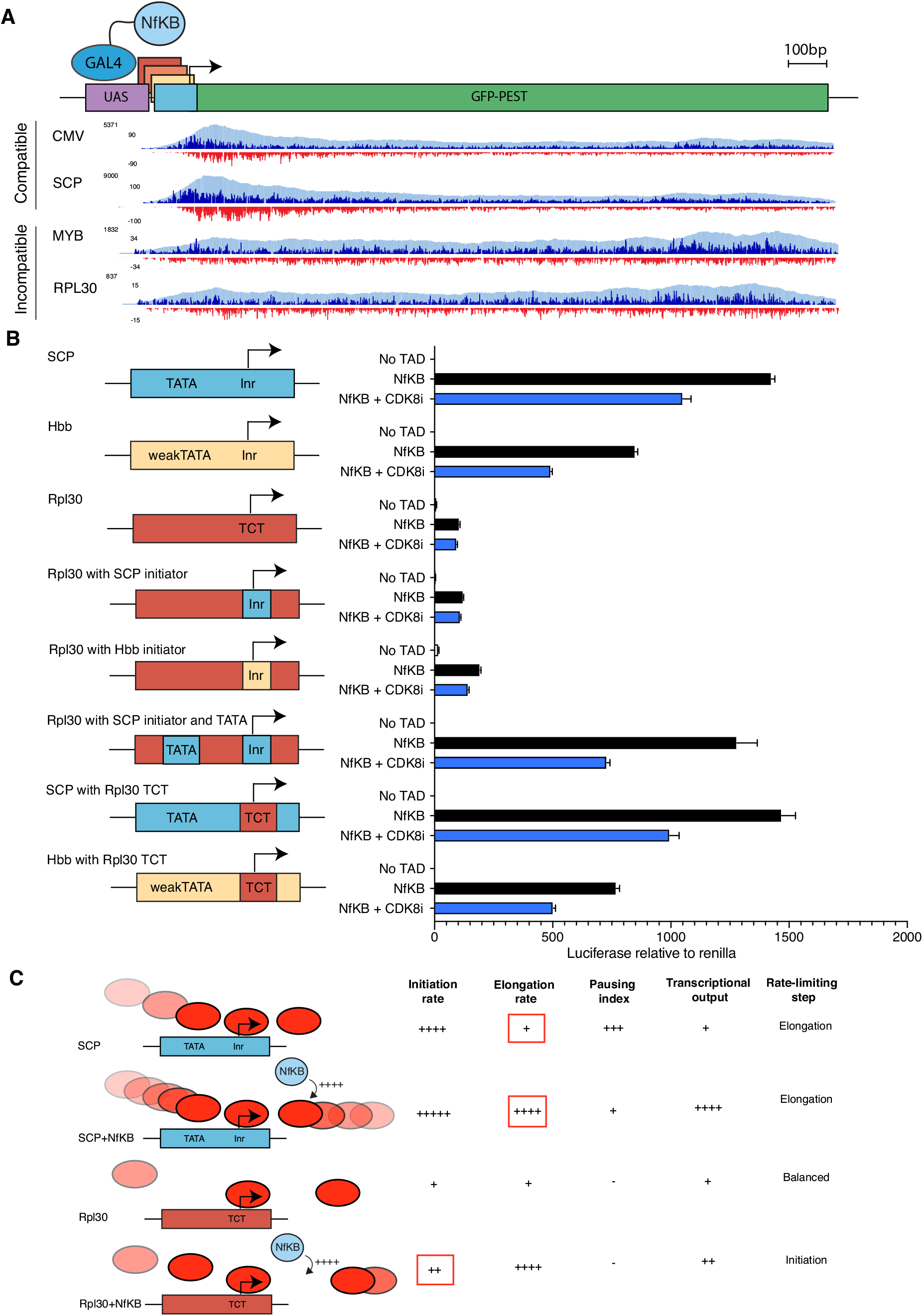
Incompatibility arises due to different rate limiting steps at certain promoters, which can be reversed through addition of a TATA box. **(A)** RNA polymerase 2 ChIP-nexus coverage across the reporter construct in compatible and incompatible promoter lines activated by Gal4-NF-kB. **(B)** Luciferase assays performed with different promoter constructs with or without Gal4-NF-kB demonstrating that adding of a TATA box restores compatibility. CDK8i was dosed for 12hrs. **(C)** Rate limiting step model of compatibility. Pause release cofactors have limited ability to activate promoters where initiation is the rate limiting step.

To investigate the contribution of each core promoter element, we swapped elements from the compatible and incompatible promoters in an attempt to alter the rate-limiting step and therefore influence cofactor-promoter compatibility (Figure 5B). Replacing the TCT motif with an Initiator motif from a TATA or TATA-like promoter did not influence transcription, demonstrating that Initiator motifs alone do not have a dominant influence on cofactor/promoter compatibility (Figure 5B). However, adding a TATA or TATA-like element and an Initiator motif into the incompatible RPL30 promoter, completely restored compatibility and markedly increased the dynamic range of gene expression (Figure 5B). Importantly, these changes to the promoter resulted in dependency on the pause release cofactor, CDK8, suggesting that pause release and elongation was now the rate-limiting step (Figure 5B). TATA boxes are known to increase the rate of transcriptional initiation by enabling efficient assembly of the pre-initiation complex (*37, 38*) and we have shown that the incompatible TCT promoters use an alternative TFIID complex that is TBP-independent (Figure 5A). This suggests that by adding a TATA box to this incompatible promoter, we altered the rate-limiting step from initiation to elongation, restored cofactor-promoter compatibility and increased the dynamic range of gene expression. Interestingly, replacing the Inr motif with a TCT element in a promoter with a TATA box had no appreciable effect on activator responsiveness (Figure 5B). Overall, our data supports a model of cofactor-promoter compatibility dictated by the presence of promoter elements that result in different rate-limiting steps for transcription (Figure 5C).

## Discussion

Understanding the rules that dictate why different genes are regulated by different transcription factors and cofactors has been a longstanding challenge for molecular biology. Here, by developing a reductionistic screening system, we have provided several key insights into the mechanisms of transcriptional regulation. Firstly, we observed that TFs rarely have dedicated cofactors that have a dominant contribution to transactivation, instead each of the specific cofactors tend to have a subtle contribution to transcription. Due to the reductionistic design of our system, we can conclude that the ability for TFs to recruit a diverse range of cofactors is predominantly determined by the sequence of their TAD, but as these TADs are unstructured, it remains unclear which precise features or motifs dictate cofactor preference. Exactly why each TF requires a unique combination of cofactors for transactivation also remains unclear. One possibility is that this type of combinatorial specificity enables TFs to integrate a large amount of information when activating their targets (*39, 40*). We provide strong experimental evidence that cofactor specificity enables different TFs to regulate distinct steps in the transactivation process. Some TFs predominantly facilitate initiation, whereas others markedly potentiate elongation. As genes are often regulated by several TFs, this combination enables kinetic synergy, as each TF activates a different rate limiting step (*41*). Kinetic synergy creates an AND logic, where only in the context of two complementary TFs does transcription proceed efficiently. Therefore, TFs could use different cofactors to create this type of logic, which may help facilitate rapid activation and reduce the prevalence of transcriptional noise (*36*).

We also extended our screens to address how changing core promoter elements influence the cofactors required for transcription. In contrast with a complex ‘lock and key’ biochemical compatibility model (*9*), our data supports a rate-limiting step model of cofactor compatibility (Figure 5C). We demonstrate that TCT promoters, which have a stable and narrow range of gene expression, are generally not constrained by the rate of pause-release and instead are restricted by initiation. In contrast, TATA gene promoters, which often regulate developmental or stress response genes that require differential expression across tissues, are constrained by pause release (*42*). Consequently, TFs such as NF-κB, which recruit cofactors that regulate pause release and elongation are less capable of increasing transcriptional output at TCT promoters. These data suggest that the pause release checkpoint may have evolved to enable a broad dynamic range of expression for genes that contain promoter elements which efficiently assemble Pol II and the PIC. In contrast, genes constrained by initiation rate, such as TCT promoters, are fundamentally less responsive to transactivation and consequently have a lower dynamic range of expression. Therefore, this suggests that promoters establish different rate-limiting steps, which are activated by different TFs, in order to achieve effective regulation of ubiquitous or inducible gene expression.

The reductionistic design we employed for all of our comparative screens has been a necessary limitation that enabled us to isolate the influence of individual variables. Whilst this approach has been effective in providing key insights into the mechanisms of transcriptional regulation, it does not capture all of the variables that contribute to transcription. Future work will need to consider the influence of other domains within TFs, the influence of regulation from distal enhancers and how different combinations of TFs contribute to endogenous mechanisms of transcriptional regulation.

---

**Figure S1).**
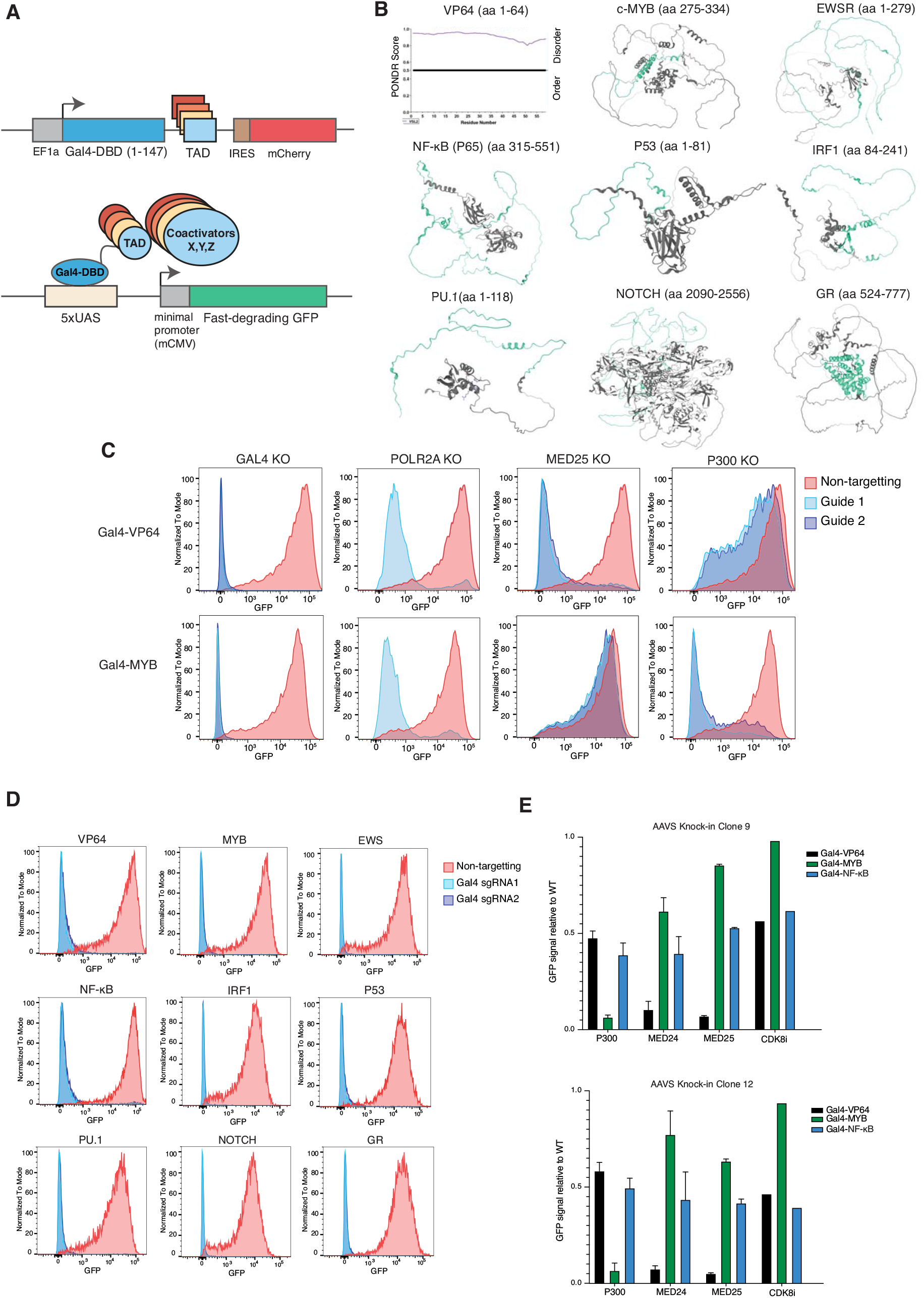
Establishing the screening system. (**A)** Schematic of the Gal4-based transactivation screening platform. TADs of interest are cloned downstream of the Gal4 DNA binding domain and recruit their respective cofactors to activate a reporter containing a fast degrading GFP. Both constructs are lentivirally integrated. **(B)** Alpha-fold predictions of the structure of the 9 transcription factors used in the CRISPR screens. The specific TAD region used is highlighted in green. VP64 does not have a reported structure and therefore PONDR analysis is displayed. **(C)** Flow cytometry analysis of the GFP signal at day 4 after infection with indicated sgRNAs in Gal4-VP64 and Gal4-MYB K562 cells. (**D)** Flow cytometry analysis of GFP signal in each of the Gal4-TAD cell lines 4 days after infection with control sgRNAs and two independent sgRNAs targeting Gal4. **(E)** Quantification of change in mean fluorescence intensity of GFP 5 days after infection with sgRNAs targeting indicated cofactors in two independent AAVS reporter knock-in clones activated by different TADs. CDK8i (Senexin A) was applied at a dose of 1uM for 24hrs.

**Figure S2).**
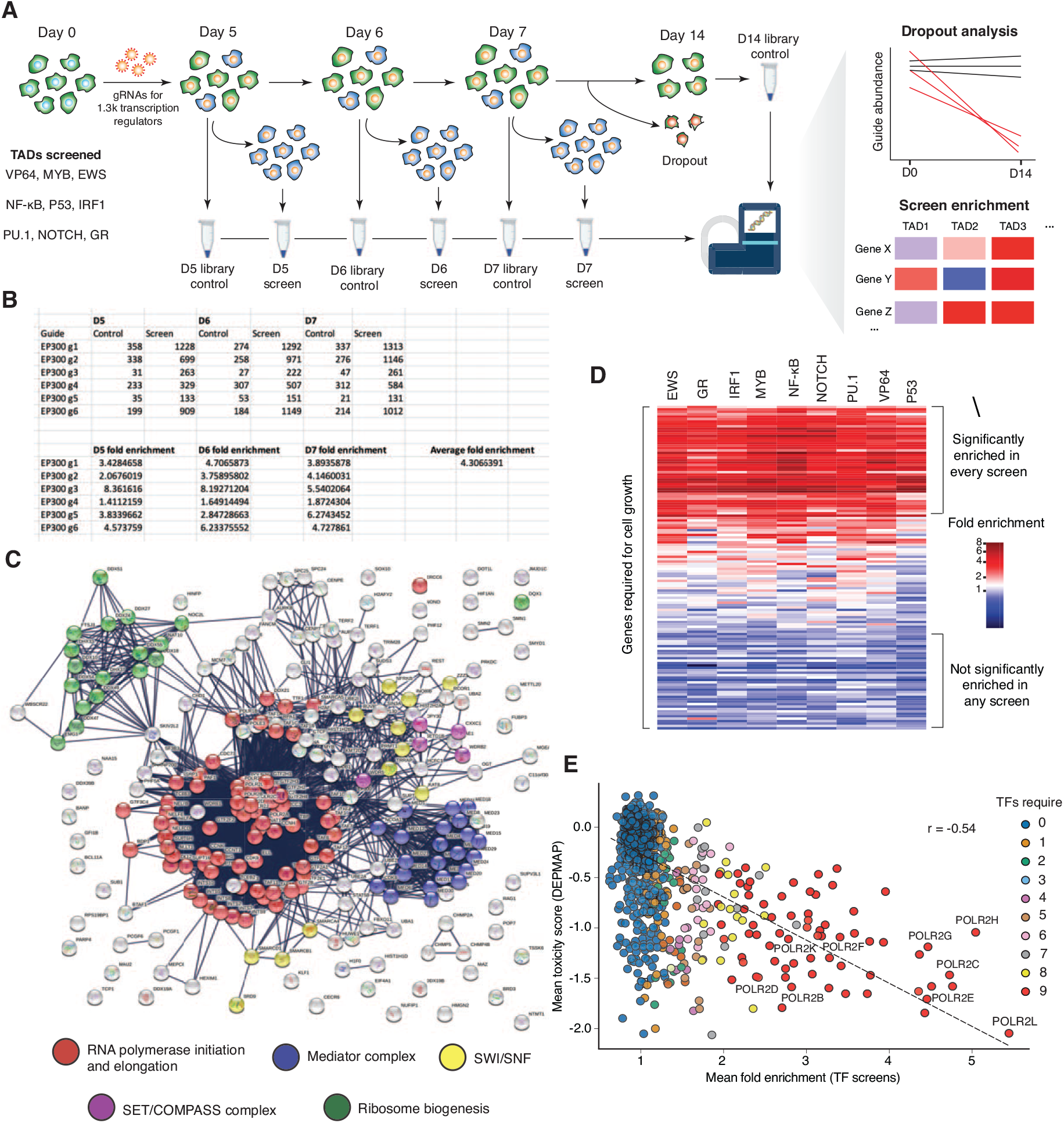
Comparative CRISPR screens performed on 9 TFs. **(A)** Schematic of the screening strategy used to identify the proteins required for each transcription factor. Dropout analysis assesses the guides depleted over time in the screen to identify genes required for cell viability. **(B)** Example of how the fold enrichment score is calculated for each specific gene in each screen. **(C)** STRING analysis on the highest stringency setting using all the genes classified as a hit for at least one of the TFs. **(D)** Fold enrichment scores from the transcriptional screens for all of the genes defined as essential for cell viability based on dropout analysis. **(E)** Correlation between the average effect of each gene in the library on transcription and the average effect of the gene on cell growth across all cell lines in the DEPMAP database. Components of the RNA polymerase 2 complex are labelled.

**Figure S3).**
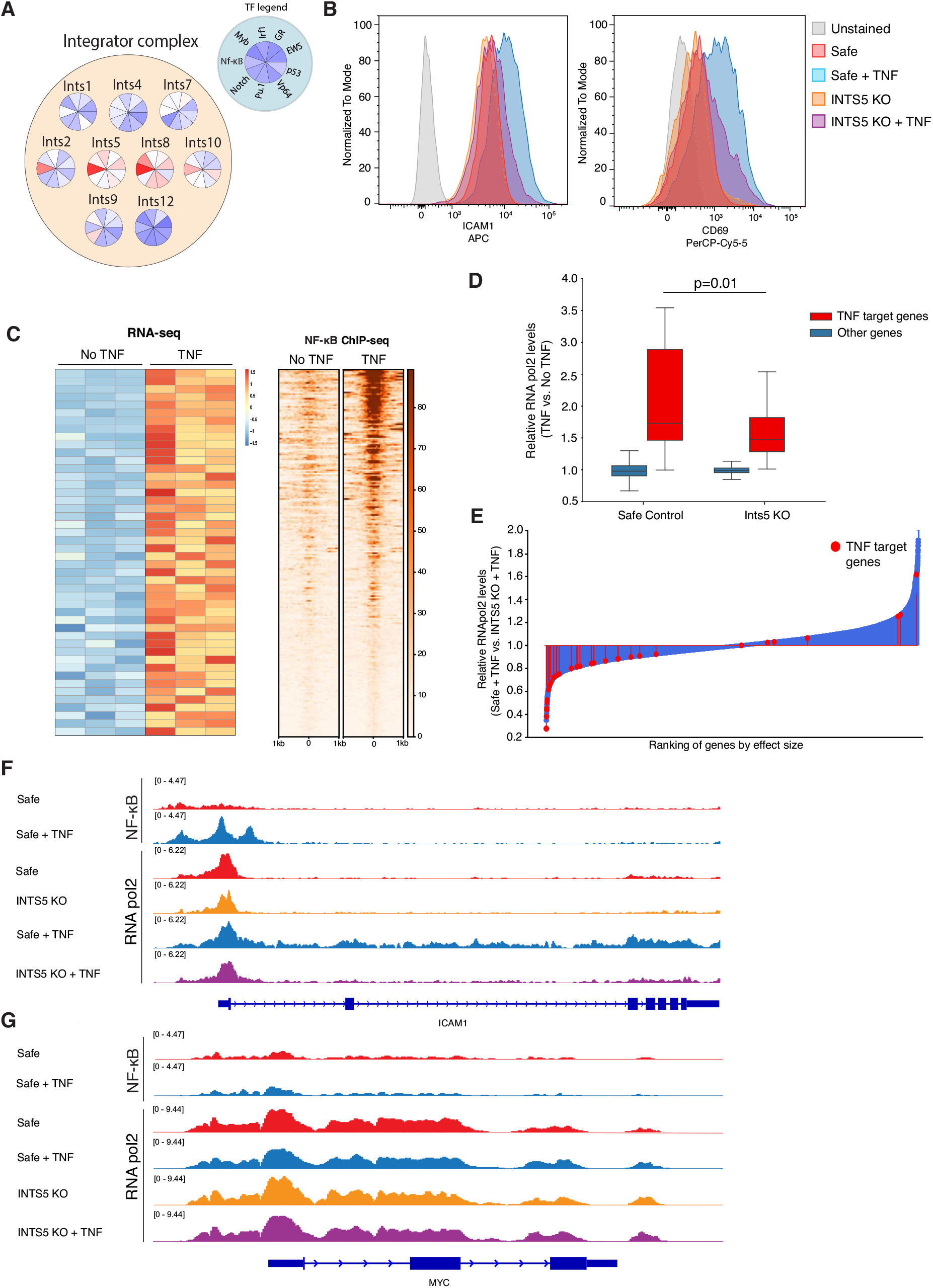
An integrator submodule containing INTS5 is required for NF-κB transactivation. **(A)** Visualisation of the requirement of Integrator subunits across each of the TFs demonstrating a disproportionate requirement for NF-κB transactivation. **(B**) Flow cytometry analysis of two TNF target genes, ICAM1 and CD69 in K562 cells with and without TNF at D4 after infection with control or INTS5 sgRNAs. **(C)** RNA-seq heatmap (left) and NF-κB ChIP-seq (right) at genes upregulated in K562 cells treated with TNF. **(D)** Quantification of the change in RNA polymerase 2 ChIP-seq signal at TNF target genes and other genes in control and INTS5 K562 cells at D4 after infection with control or INTS5 sgRNAs. **(E)** Waterfall plot of change RNA polymerase 2 levels in INTS5 KO K562 cells treated with TNF relative to control K562 cells treated with TNF. TNF target genes are highlighted in red. **(F)** IGV snapshot of NF-κB or RNA polymerase 2 occupancy at ICAM1 in control and INTS5 KO K562 cells, with and without TNF. **(G)** IGV snapshot of NF-κB or RNA polymerase 2 occupancy at MYC in control and INTS5 KO K562 cells, with and without TNF.

**Figure S4).**
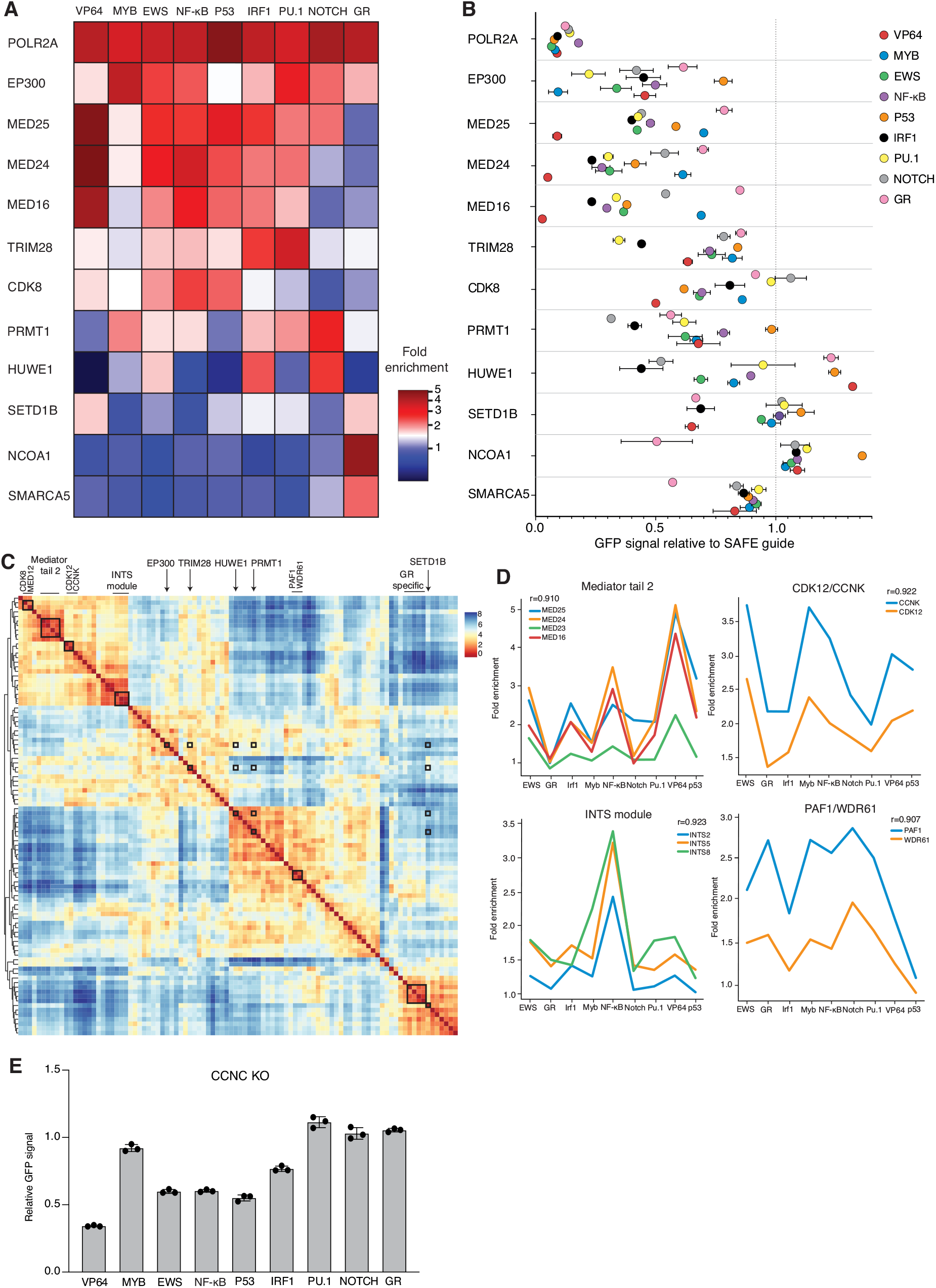
Global analysis of TAD CRISPR screens. **(A)** Heatmap of fold enrichment score from the screen data for selected set of genes. * indicates gene classified as significant hit. **(B)** Validation of screen data through flow cytometry based quantification of GFP signal. Quantification value is based on M.F.I and is compared to a SAFE guide control at D5 after infection with relevant sgRNAs. Most genes displayed were validated with at least 2 sgRNAs, error bars reflect S.E.M between two sgRNAs. **(C)** Correlation matrix of the top 100 most heterogeneous regulators. Correlation score based on Euclidean distance. Clear co-dependent clusters are highlighted. Other independently validated heterogeneous hits that do not display correlated patterns of requirement are also highlighted. **(D)** Fold enrichment scores for co-dependent gene clusters. r value reflects average correlation between patterns of enrichment. **(E)** Quantification of the effect of Cyclin C (CCNC) KO on transactivation by each of the transcription factors, based on GFP signal. Error bars reflect S.E.M of 3 different sgRNAs.

**Figure S5).**
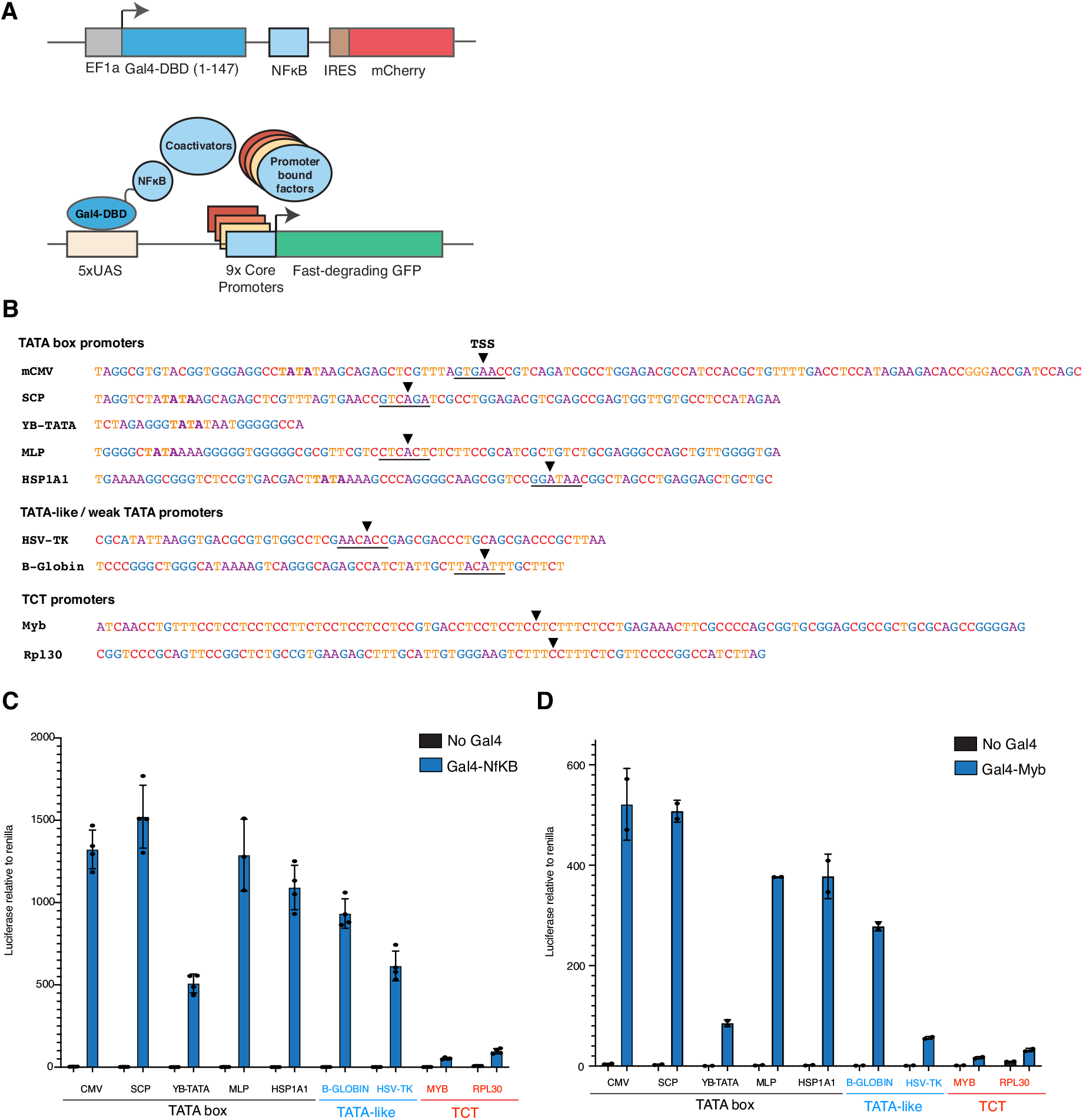
Comparative screens to assess the effect of the core promoter on the mechanisms of transactivation. (**A)** Schematic of the Gal4-based promoter screening platform. Promoters of interest are cloned downstream of the UAS and are activated by NF-κB and its associated cofactors. Both constructs are lentivirally integrated. **(B)** Sequences of the different promoters screened. The TSS is indicated with an arrow. TATA box sequences are shown in bold and the Initiator sequence is underlined. **(C)** Luciferase assays performed with different promoter constructs activated by Gal4-NF-κB. **(D)** Luciferase assays performed with different promoter constructs activated by Gal4-Myb.

## Methods

### Cell culture

A clonal K562 Cas9 cell line was generated previously to ensure high efficiency CRISPR editing (*1*). K562 cells were cultured in RPMI-1640 supplemented with 20% FCS, streptomycin (100ug/ml), penicillin (100 units/ml) and glutamax, under standard culture conditions (5% CO2, 37°C). HEK293ET cells were grown in DMEM supplemented with 10% FCS, streptomycin (100ug/ml), penicillin (100 units/ml) and glutamax, under standard culture conditions (5% CO2, 37°C). All cell lines were subjected to regular mycoplasma testing and underwent short tandem repeat (STR) profiling.

### Lentivirus production and transduction

Lentivirus was prepared by transfecting HEK293ET cells with plasmid:pVSVg:psPAX2 plasmids in a 3:2:1 ratio using PEI reagent. The viral supernatant was collected 48-72hrs following transfection, filtered through a 0.45 μm filter, and added to cells.

### Drug treatment

Senexin A (CDK8i) (Selleckchem) was dosed at 1uM for 24hrs. Gal4-GR cells were dosed with 1uM Dexamethasone (Sigma) to induce GR activity. TNF-a treatment was performed for 16hrs at 25ng/ml.

### Flow cytometry analyses

Flow cytometry analyses were performed on the LSRFortessa X-20 flow cytometer (BD Biosciences). Data were analyzed with FlowJo software (Tree Star). Cell sorting was performed on the FACS FACSAria Fusion flow sorter (BD Biosciences).

### Cloning of screening system

pKLV-U6gRNA(BbsI)-Puro2ABFP vector was used as the base vector for cloning the lentiviral Gal4-TAD vector. The entire gRNA, Puromycin and BFP regions of the plasmid were removed and replaced by the Gal4-DBD together with an IgA linker through standard cut and paste cloning. The EF1a, IRES and Cherry were then introduced sequentially. The vector was designed for simple cut and paste replacement of the TAD region downstream of the Gal4-DBD with alternative TADs which were PCR’d out of cDNA expression vectors obtained from Addgene. Primers used are listed in Data S5. The base for the reporter construct was obtained from Addgene (#79199). The original vector is a lentiviral vector that includes the 5xUAS upstream of a minimal CMV promoter. Downstream of this promoter, the Turbo-GFP-PEST was subcloned from another Addgene vector (#67180). To produce different promoter reporters, promoter regions were obtained from previous publications or from the eukaryotic promoter database. For endogenous promoters, a region from ~ −80bps to +20bps of the reported TSS was used. Oligos corresponding to these regions were produced, annealed and cloned into the 5xUAS-mCMV-Turbo-GFP-PEST construct, replacing the mCMV promoter.

### Generation of Gal4-TAD and promoter reporter cell lines

For TAD screens, the lentiviral 5xUAS mCMV Turbo-GFP-PEST reporter was introduced into clonal K562-Cas9-Blastycidine cell line at a high MOI to minimise locus specific effects and to ensure robust detection of GFP signal upon activation by the TADs. Into this reporter line, each of the Gal4-TAD constructs was introduced by lentiviral integration. For promoter screens, the lentiviral 5xUAS Turbo-GFP-PEST reporter with variable promoters were integrated into a clonal K562-Cas9-Blastycidine cell line at high MOI. For both sets of cell lines, GFP positive cells were sorted from each cell line, until a pure and stable GFP population was obtained.

### sgRNA design and cloning

sgRNAs were designed using the IDT CRISPR design tool or were obtained from the sequences of guides in the pooled guide library. sgRNAs were cloned into the pKLV-U6gRNA(BbsI)-Puro2ABFP vector using standard golden gate cloning.

### sgRNA and primer sequences

The sequences for sgRNA sequences and primers are included in Data S5.

### Validation and quantification of screen hits

All experiments validating and quantifying the effect of cofactor knockout were performed by quantifying the mean fluorescence intensity (MFI) of the GFP signal at D5 after sgRNA infection. Each TAD line was infected with the same batch of virus in parallel to minimise technical variation between the quantification. Most genes were validated with at least 2 guides and at least 2 biological replicates.

### HDR mediated AAVS knock-in

The 5xUAS, mCMV promoter and Turbo-GFP-PEST were cloned reporter by Gibson assembly into the pMK232 (CMV-OsTIR1-Puro), which contained homology arms for the AAVS locus. The sgRNA targeting the AAVS was introduced into the px330-mcherry vector, which contains both Cas9 and the gRNA. The AAVS-reporter repair template and the px330-Cas9-AAVS gRNA vector was electroporated into K562 cells using the Neon Transfection system (Thermo Fischer Scientific) with settings optimised for K562 cells. Single cell clones were sorted 5 days after transfection and grown out for 2 weeks to obtain sufficient cells. AAVS knock-in clones were identified by In-Out PCR and Sanger sequencing.

### Guide library design, generation and cloning

To assess the requirement of transcriptional regulators, a bespoke library of gRNAs that targets over 1137 known chromatin and transcriptional regulators was designed (Data S2). The library was designed through a combination of searches for genes containing domains known to be enriched in transcriptional regulators and manual curation. Each gene was targeted with 6 independent gRNAs. As controls, the library also contains a large number of guides targeting safe regions and guides that do not target any genomic locus. The total library contains 7239 gRNAs. The oligo pool was synthesized by CustomArray (Genescript). The sgRNA pool was PCR amplified and pot cloned into the pKLV-U6gRNA(BbsI)-Puro2ABFP vector using standard cut and paste cloning. The ligated product was electroporated into Electrocompetent cells (Lucigen) and grown in liquid culture overnight at 37 degrees before being extracted by Maxiprep. Low skewing of the plasmid was confirmed by sequencing of the cloned plasmid pool.

### Comparative CRISPR screens

Prior to beginning the screens, reporter cell lines were sorted to obtain a pure GFP positive population. Sufficient cells were used to maintain 1000-fold representation at all stages of the screening process. The cells were transduced with an appropriate volume of viral supernatant to ensure only a single guide was present in most cells (MOI=0.3). At day 5, 6 and 7 day after guide infection, guide positive, GFP negative cells (< 25% of the MFI of the entire population) were sorted. Guide positive cells were also sorted as a library control at each time point to provide a library control reference to calculate enrichment. Four of the Gal4-TAD cell lines from the comparative CRISPR screens were also maintained until day 14 after guide infection to test for genes required for cell growth. These four TAD lines were used as independent replicates for the dropout analysis. Genomic DNA was extracted using Monarch^®^ Genomic DNA purification kit (New England Biolabs), according to the manufacturer’s instructions. PCR was conducted to maintain guide representation, using Q5^®^ High Fidelity DNA Polymerase (New England Biolabs). PCR products were pooled and sequenced on the NextSeq500 using 75bp paired-end chemistry.

### Analysis of comparative CRISPR screens

The sequence reads were trimmed to remove the constant portion of the sgRNA sequences with cutadapt, then mapped to the reference sgRNA library with Bowtie2 (*2*). After filtering to remove multi-mapping reads, the read counts were computed for each sgRNA. After obtaining guide counts for all samples, a series of filtering and processing steps were performed to calculate mean fold enrichment values for each gene in each screen. Firstly, guides that were very lowly represented (below 2.5^th^ percentile) were filtered from the analysis, since their low representation caused extreme fold change values. The counts were then normalised to sequencing depth before calculating a fold enrichment score for each guide by dividing the counts for each gene in the screen samples by the counts in the library control. This resulted in a total of 18 individual fold enrichment scores for each gene (6 guides, 3 timepoints). Given that we still observed some examples of extreme fold change values that would obscure the calculation of a mean fold change value for each gene, we filtered any guides that had a fold change below 0.1 or greater than 10. Using the remaining guides, we were able to calculate a fold enrichment score for each gene. Further outliers were removed by filtering guides that were more than 4-fold away from the mean fold change value.

Using this filtered guide list, we calculated a final fold enrichment score for each gene in each screen. In order to calculate what fold enrichment score should be considered statistically significant, a permutation test was performed for each screen. The permutation test shows what fold change distribution would be expected if you randomly sampled fold enrichment scores from guides in the data. Specifically, 6 guides were randomly sampled from each timepoint providing a vector of 18 values. The mean fold enrichment was calculated across these 18 values. Random sampling was performed 10000 times to produce a random sampling distribution. Genes that had a mean fold change and at least 1/3^rd^ of the values (6/18) above the 95^th^ percentile of this distribution were considered significant.

To identify heterogeneous regulators, a heterogeneity score was calculated for each gene. The heterogeneity score reflects the standard deviation of a gene across all 9 screens divided by its mean fold enrichment score. A correlation matrix of the ~100 top heterogeneous regulators was based on the similarity of patterns in requirement using Euclidean distance. To define which genes affected cell growth, MAGeCK analysis was performed using the D14 timepoint from 4 of the TAD screens, comparing them to the plasmid as the D0 reference (*3*). Any genes with an adjusted p-value below 0.05 were considered significant.

### Cofactor KO ChIP-seq and RNA-seq experiments

sgRNAs targeting MED12, MED14, MED16, MED24, MED25, CDK8 and CCNC were lentivirally introduced into a K562-Cas9 clone. Guide efficiency was validated through flow cytometry analysis of phenotypic effects of guide knockout. Cells were grown for 4 days after infection at which timepoint they were harvested for ChIP-seq.

### Chromatin immunoprecipitation (ChIP)

For each ChIP, at least 20 million cells were crosslinked for 15 mins with 1% formaldehyde. Crosslinked material was sonicated to approximately 200-1000bp using the Covaris Ultrasonicator e220. Sonicated material was incubated overnight with each antibody, then incubated for 3hrs with Protein A magnetic beads. Beads were washed with low and high salt wash buffers, LiCl buffer and TE, before being eluted and de-crosslinked overnight. DNA was purified using Qiagen Minelute columns. All ChIP antibodies were used at ~10ug per IP. Sequencing libraries were prepared from eluted DNA using Rubicon ThruPLEX DNA-seq kit. Libraries were size selected between 200-500bps and sequenced on the NextSeq500 using the 75bp single-end chemistry. The following antibodies were used for ChIP analysis: Mouse anti-RNA polymerase II antibody clone CTD4H8 (Merck Millipore, 05-623), Rabbit anti-NF-kB p65 antibody clone D14E12 (Cell Signalling, 8242).

### ChIP-seq analysis

Reads were aligned to the human genome (GRCh38) with Bowtie2 (*2*). Duplicate reads and reads mapping to blacklist regions or the mitochondria were removed. ChIP-seq coverage across selected genomic regions was calculated with BEDtools (*4*).

### Mediator KO RNA Polymerase 2 ChIP-seq analysis

After alignment and depth normalisation, the coverage across genes was calculated using BEDtools. To define which genes are Mediator dependent, we took the 10000 genes with the most RNA polymerase 2 signal across the gene and identified genes with at least 30% reduction in total RNA polymerase 2 signal in the MED14 KO.

### ChlP-nexus

ChIP-nexus was performed as described previously (*5–7*). Briefly, the immunoprecipitation and washes were performed using the same conditions as our ChIP protocol. Upon competition of these steps, the DNA was end-repaired, A-tailed, adaptors ligated, exonuclease treated, circularized on Dynabeads as described previously (*5–7*). DNA was then eluted from the beads and PCR was performed to produce sequencing ready libraries. For ChIP-nexus performed on the Gal4-TAD lines, DNA was sequenced on the NextSeq500 using the 75bp single-end chemistry. For ChIP-nexus on the promoter lines, DNA was sequenced on the NextSeq500 using 75bp single-end chemistry, run to produce paired-end 37bp reads.

### Luciferase assays

To generate the reporter constructs for the luciferase assays, promoters of interest were cloned into the pGL4.35 (luc2p/9xgal4uas/hygro) luciferase construct (Promega, E1370) downstream of the 9xUAS site. Luciferase constructs were introduced into HEK293T cells by transient transfection using PEI. The luciferase construct of interest was co-transfected with or without the relevant Gal4-TAD and the pRL Renilla control (Promega, E2261). Cells were harvested 48hrs after transfection. Luciferase signal and Renilla signal were analysed using the Dual-Luciferase reporter system (Promega, E1910) using the Cytation 3 plate-reader (BioTek).

## Acknowledgements

The authors would like to acknowledge all members of the Dawson Lab for their support and intellectual input throughout the project. We would also like to acknowledge the Peter MacCallum Cancer Centre Flow Cytometry and Genomics core facilities for their assistance with the research.

## Funding

This research in the Dawson lab was supported by a Cancer Council Victoria Postdoctoral fellowship (C.C.B), NHMRC Investigator Grant (1196749, M.A.D.), Cancer Council Victoria Dunlop Fellowship (M.A.D), Howard Hughes Medical Institute international research scholarship (55008729, M.A.D) and ARC project grant (DP220103927).

## Contributions

C.C.B, O.G and M.A.D designed the research and interpreted the data. C.C.B and M.A.D supervised the research and co-wrote the manuscript with helpful input from all the authors. C.C.B performed the experiments with assistance from L.S. C-S.A and O.G. L.T performed the bioinformatic analysis with assistance from E.Y.N.

## Competing interests

M.A.D. has been a member of advisory boards for GSK, CTX CRC, Storm Therapeutics, Celgene, and Cambridge Epigenetix and receives research funding from Pfizer.

## References

1. S. A. Lambert, A. Jolma, L. F. Campitelli, P. K. Das, Y. Yin, M. Albu, X. Chen, J. Taipale, T. R. Hughes, M. T. Weirauch, The Human Transcription Factors. Cell. 172, 650–665 (2018).

2. M. Ptashne, A. Gann, Transcriptional activation by recruitment. Nature. 386, 569–577 (1997).

3. R. G. Roeder, Transcriptional regulation and the role of diverse coactivators in animal cells. FEBS Lett. 579, 909–915 (2005).

4. R. Donczew, L. Warfield, D. Pacheco, A. Erijman, S. Hahn, Two roles for the yeast transcription coactivator SAGA and a set of genes redundantly regulated by TFIID and SAGA. Elife. 9, 1–27 (2020).

5. F. Reiter, S. Wienerroither, A. Stark, Combinatorial function of transcription factors and cofactors. Curr. Opin. Genet. Dev. 43, 73–81 (2017).

6. F. Nemčko, A. Stark, Proteome-scale identification of transcriptional activators in human cells. Mol. Cell. 82, 497–499 (2022).

7. D. T. Bergman, T. R. Jones, V. Liu, J. Ray, E. Jagoda, L. Siraj, H. Y. Kang, J. Nasser, M. Kane, A. Rios, T. H. Nguyen, S. R. Grossman, C. P. Fulco, E. S. Lander, J. M. Engreitz, Compatibility rules of human enhancer and promoter sequences. Nature. 106, 1–22 (2022).

8. M. Martinez-Ara, F. Comoglio, J. van Arensbergen, B. van Steensel, Mol. Cell, in press, doi:10.1016/j.molcel.2022.04.009.

9. J. Van Arensbergen, B. Van Steensel, H. J. Bussemaker, In search of the determinants of enhancer – promoter interaction specificity. Trends Cell Biol. 24, 695–702 (2014).

10. C. C. Galouzis, E. E. M. Furlong, Regulating specificity in enhancer-promoter communication. Curr. Opin. Cell Biol. 75, 102065 (2022).

11. M. Levine, C. Cattoglio, R. Tjian, Looping Back to Leap Forward: Transcription Enters a New Era. Cell. 157, 13–25 (2014).

12. J. L. Schmid-Burgk, K. Höning, T. S. Ebert, V. Hornung, CRISPaint allows modular base-specific gene tagging using a ligase-4-dependent mechanism. Nat. Commun. 7, 12338 (2016).

13. D. R. Pattabiraman, C. McGirr, K. Shakhbazov, V. Barbier, K. Krishnan, P. Mukhopadhyay, P. Hawthorne, A. Trezise, J. Ding, S. M. Grimmond, P. Papathanasiou, W. S. Alexander, A. C. Perkins, J. P. Levesque, I. G. Winkler, T. J. Gonda, Interaction of c-Myb with p300 is required for the induction of acute myeloid leukemia (AML) by human AML oncogenes. Blood. 123, 2682–2690 (2014).

14. E. Vojnic, A. Mourão, M. Seizl, B. Simon, L. Wenzeck, L. Larivière, S. Baumli, K. Baumgart, M. Meisterernst, M. Sattler, P. Cramer, Structure and VP16 binding of the Mediator Med25 activator interaction domain. Nat. Struct. Mol. Biol. 18, 404–409 (2011).

15. M. L. Sandberg, S. E. Sutton, M. T. Pletcher, T. Wiltshire, L. M. Tarantino, J. B. Hogenesch, M. P. Cooke, c-Myb and p300 regulate hematopoietic stem cell proliferation and differentiation. Dev. Cell. 8, 153–166 (2005).

16. Y. Xu, J. P. Milazzo, T. D. D. Somerville, Y. Tarumoto, Y.-H. Huang, E. L. Ostrander, J. E. Wilkinson, G. A. Challen, C. R. Vakoc, A TFIID-SAGA Perturbation that Targets MYB and Suppresses Acute Myeloid Leukemia. Cancer Cell. 33, 13–28.e8 (2018).

17. A. J. Donner, S. Szostek, J. M. Hoover, J. M. Espinosa, CDK8 Is a Stimulus-Specific Positive Coregulator of p53 Target Genes. Mol. Cell. 27, 121–133 (2007).

18. C. Y. Chung, Z. Sun, G. Mullokandov, A. Bosch, Z. A. Qadeer, E. Cihan, Z. Rapp, R. Parsons, J. A. Aguirre-Ghiso, E. F. Farias, B. D. Brown, A. Gaspar-Maia, E. Bernstein, Cbx8 Acts Non-canonically with Wdr5 to Promote Mammary Tumorigenesis. Cell Rep. 16, 472–486 (2016).

19. H. Zheng, Y. Qi, S. Hu, X. Cao, C. Xu, Z. Yin, X. Chen, Y. Li, W. Liu, J. Li, J. Wang, G. Wei, K. Liang, F. X. Chen, Y. Xu, Identification of Integrator-PP2A complex (INTAC), an RNA polymerase II phosphatase. Science. 370 (2020), doi:10.1126/science.abb5872.

20. L. El Khattabi, H. Zhao, J. Kalchschmidt, N. Young, S. Jung, P. Van Blerkom, P. Kieffer-Kwon, K.-R. Kieffer-Kwon, S. Park, X. Wang, J. Krebs, S. Tripathi, N. Sakabe, D. R. Sobreira, S.-C. Huang, S. S. P. Rao, N. Pruett, D. Chauss, E. Sadler, A. Lopez, M. A. Nóbrega, E. L. Aiden, F. J. Asturias, R. Casellas, A Pliable Mediator Acts as a Functional Rather Than an Architectural Bridge between Promoters and Enhancers. Cell. 178, 1145–1158 (2019).

21. R. Abdella, A. Talyzina, S. Chen, C. J. Inouye, R. Tjian, Y. He, Structure of the human Mediator-bound transcription preinitiation complex. Science. 372, 52–56 (2021).

22. W. Shao, J. Zeitlinger, Paused RNA polymerase II inhibits new transcriptional initiation. Nat. Genet. 49, 1045–1051 (2017).

23. Y. Aoi, E. R. Smith, A. P. Shah, E. J. Rendleman, S. A. Marshall, A. R. Woodfin, F. X. Chen, R. Shiekhattar, A. Shilatifard, NELF Regulates a Promoter-Proximal Step Distinct from RNA Pol II Pause-Release. Mol. Cell. 78, 261–274.e5 (2020).

24. K. M. Leach, K. F. Vieira, S. H. Lee Kang, A. Aslanian, M. Teichmann, R. G. Roeder, J. Bungert, Characterization of the human β-globin downstream promoter region. Nucleic Acids Res. 31, 1292–1301 (2003).

25. J. J. Stewart, J. A. Fischbeck, X. Chen, L. A. Stargell, Non-optimal TATA elements exhibit diverse mechanistic consequences. J. Biol. Chem. 281, 22665–22673 (2006).

26. J. J. Stewart, L. A. Stargell, The Stability of the TFIIA-TBP-DNA Complex Is Dependent on the Sequence of the TATAAA Element. J. Biol. Chem. 276, 30078–30084 (2001).

27. J. T. Kadonaga, Perspectives on the RNA polymerase II core promoter. Wiley Interdiscip. Rev. Dev. Biol. 1, 40–51 (2012).

28. J. M. Morachis, C. M. Murawsky, B. M. Emerson, Regulation of the p53 transcriptional response by structurally diverse core promoters. Genes Dev. 24, 135–147 (2010).

29. T. J. Parry, J. W. M. Theisen, J. Y. Hsu, Y. L. Wang, D. L. Corcoran, M. Eustice, U. Ohler, J. T. Kadonaga, The TCT motif, a key component of an RNA polymerase II transcription system for the translational machinery. Genes Dev. 24, 2013–2018 (2010).

30. V. Haberle, C. D. Arnold, M. Pagani, M. Rath, K. Schernhuber, A. Stark, Transcriptional cofactors display specificity for distinct types of core promoters. Nature. 570, 122–126 (2019).

31. A. B. Patel, R. K. Louder, B. J. Greber, S. Grünberg, J. Luo, J. Fang, Y. Liu, J. Ranish, S. Hahn, E. Nogales, Structure of human TFIID and mechanism of TBP loading onto promoter DNA. Science. 362, eaau8872 (2018).

32. N. Petrenko, Y. Jin, L. Dong, K. H. Wong, K. Struhl, Requirements for RNA polymerase II preinitiation complex formation in vivo. Elife. 8, 1–19 (2019).

33. R. K. Louder, Y. He, J. R. López-blanco, J. Fang, P. Chacón, E. Nogales, Structure of promoter-bound TFIID and model of human pre-initiation complex assembly. Nature (2016), doi:10.1038/nature17394.

34. H. Kwak, N. J. Fuda, L. J. Core, J. T. Lis, Precise Maps of RNA Polymerase Reveal How Promoters Direct Initiation and Pausing. Science. 339, 950–953 (2013).

35. D. A. Gilchrist, G. Dos Santos, D. C. Fargo, B. Xie, Y. Gao, L. Li, K. Adelman, Pausing of RNA polymerase II disrupts DNA-specified nucleosome organization to enable precise gene regulation. Cell. 143, 540–551 (2010).

36. L. Core, K. Adelman, Promoter-proximal pausing of RNA polymerase II: A nexus of gene regulation. Genes Dev. 33, 960–982 (2019).

37. B. C. Hoopes, J. F. LeBlanc, D. K. Hawley, Contributions of the TATA box sequence to rate-limiting steps in transcription initiation by RNA polymerase II. J. Mol. Biol. 277, 1015–1031 (1998).

38. D. Yean, J. Gralla, Transcription reinitiation rate: a special role for the TATA box. Mol. Cell. Biol. 17, 3809–3816 (1997).

39. H. E. Klumpe, M. A. Langley, J. M. Linton, C. J. Su, Y. E. Antebi, M. B. Elowitz, The context-dependent, combinatorial logic of BMP signaling. Cell Syst. 13, 388–407.e10 (2022).

40. C. J. Su, A. Murugan, J. M. Linton, A. Yeluri, J. Bois, H. Klumpe, M. A. Langley, Y. E. Antebi, M. B. Elowitz, Ligand-receptor promiscuity enables cellular addressing. Cell Syst. 13, 408–425.e12 (2022).

41. D. Herschlag, F. B. Johnson, Synergism in transcriptional activation: a kinetic view. Genes Dev. 7, 173–179 (1993).

42. C. Yang, E. Bolotin, T. Jiang, F. M. Sladek, E. Martinez, Prevalence of the initiator over the TATA box in human and yeast genes and identification of DNA motifs enriched in human TATA-less core promoters. Gene. 389, 52–65 (2007).

